# TRβ Agonism Induces Tumor Suppression and Enhances Drug Efficacy in Anaplastic Thyroid Cancer in Female Mice

**DOI:** 10.1101/2021.06.09.447689

**Authors:** Noelle E. Gillis, Lauren M. Cozzens, Emily R. Wilson, Noah M. Smith, Jennifer A. Tomczak, Eric L. Bolf, Frances E. Carr

**Affiliations:** Department of Pharmacology, Larner College of Medicine, University of Vermont, Burlington, Vermont; University of Vermont Cancer Center, University of Vermont, Burlington, Vermont; Current address: Masonic Cancer Center, University of Minnesota, Minneapolis, Minnesota; Current address: University of Colorado, Anschutz Medical Campus, Aurora, Colorado; Current address: University of Utah, Salt Lake City, Utah

**Keywords:** GC-1, Sobetirome, thyroid hormone receptor, anaplastic thyroid cancer

## Abstract

Thyroid hormone receptor beta (TRβ) is a recognized tumor suppressor in numerous solid cancers. The molecular signaling of TRβ has been elucidated in several cancer types through re-expression models. Remarkably, the potential impact of selective activation of endogenous TRβ on tumor progression remains largely unexplored. We used cell-based and *in vivo* assays to evaluate the effects of the TRβ agonist Sobetirome (GC-1) on a particularly aggressive and dedifferentiated cancer, anaplastic thyroid cancer (ATC). Here we report that GC-1 reduced the tumorigenic phenotype, decreased cancer stem-like cell populations, and induced re-differentiation of the ATC cell lines with different mutational backgrounds. Of note, this selective activation of TRβ amplified the effects of therapeutic agents in blunting the aggressive cell phenotype and stem-cell growth. In xenograft assays, GC-1 alone inhibited tumor growth and was as effective as the kinase inhibitor, Sorafenib. These results indicate that selective activation of TRβ not only induces a tumor suppression program *de novo* but enhances the effectiveness of anti-cancer agents revealing potential novel combination therapies for ATC and other aggressive solid tumors.

## INTRODUCTION

There is compelling evidence that the nuclear thyroid hormone receptor beta (TRβ) blunts tumor progression, invasion, and metastases in multiple solid tumors including thyroid, breast, colon and other cancers^1^. Whereas diminished expression and/or function of TRβ is often characteristic of advanced and aggressive cancers, cell-based and animal studies reveal that liganded re-expressed TRβ induces apoptosis, reduces an aggressive phenotype, decreases cancer stem cell populations, and slows tumor growth through modulation of a complex transcriptional network^2–9^. Transcriptional regulation by TRβ is critical for its function as a tumor suppressor because it also acts as both a signal transducer and facilitator of long-term epigenetic programming for maintenance of cell identity. Our recent studies revealed that TRβ in the presence of triiodothyronine (T_3_) induced a tumor-suppressive transcriptomic program and cell redifferentiation in aggressive thyroid and breast cancer cells^2,7^. We identified several key mechanisms by which activation of TRβ could reduce tumor progression, implicating thyroid hormone signaling as a therapeutic opportunity in advanced cancers. Recent studies in breast ^8^, colon^10^, and liver^5^ cancers also emphasize this untapped therapeutic potential^11,12^. Despite the potential clinical benefits of TRβ tumor suppressor activity implicated by numerous cell-based and animal studies, TRβ selective agonists remain largely untested as cancer therapeutics as single agents or in combination with other treatments. Novel treatments are a critical need in the context of aggressive poorly differentiated tumors, such as anaplastic thyroid cancer (ATC), for which there are limited targeted therapies and thus far, no curative options.

As noted in ATC and other endocrine-related cancers, activation of phosphatidylinositol 3-kinase (PI3K)–Akt–mammalian target of rapamycin (mTOR) pathway (PI3K-Akt) and mitogen-activated protein kinase (MAPK) signaling are recognized drivers of thyroid oncogenesis and blocking this activity has been a therapeutic focus. Yet inhibitors of the major components of these and other signaling cascades rarely provide a durable response as resistance quickly develops^13^. Current molecular diagnostics provide a rationale for intervention with combination therapies but thus far have not yielded significantly improved responses and potential use of thyromimetics not systematically evaluated^14 13,15,16^. The therapuetic use of non-isoform selective thyromimetics is problematic due to the side-effects of hyperthyroidism and the potential for adverse cardiovascular events that result from aberrant TRα activity.

To subvert these negative side effects, TRβ-selective thyromimetics have been developed^17^. Several of these drugs have been shown to be effective for treatment of metabolic and neurodegenerative disorders in clinical trials^17,18^. The TRβ agonist Sobetirome (GC-1) has been extensively characterized and shown to preferentially bind to and activate TRβ rather than TRα^19–22^. Thus, given our prior work demonstrating that restoration of TRβ expression reduces the aggressive ATC phenotype^2,7,23^, we here tested the possibility that selective activation of endogenous TRβ with GC-1 might reduce the tumor cell phenotype and inhibit tumor growth. In ATC cells with diverse genetic backgrounds ^24^ and varying endogenous TRβ levels, GC-1 alone decreased the aggressive phenotype, reduced cancer stem cell growth and increased the effectiveness of MAPK, PI3K, and cell cycle inhibitors. GC-1 significantly increased expression of thyroid-specific genes, inducing re-differentiation in ATC cells.

Importantly, we observed an increase in NIS protein and cellular iodide uptake, implicating potential restoration of functionality. In a xenograft model, GC-1 reduced tumor growth as effectively as a MAPK pathway inhibitor, Sorafenib. The combination of GC-1 and Sorafenib maximally suppressed tumor growth. These observations establish the foundation that activation of endogenous TRβ with isoform-selective agonists may be an effective and practical adjuvant therapeutic strategy.

## MATERIALS AND METHODS

### Culture of thyroid cell lines

ATC cell lines were cultured in RPMI 1640 growth media with L-glutamine (300 mg/L), sodium pyruvate and nonessential amino acids (1%) (Corning), supplemented with 10% fetal bovine serum (Peak Serum) and penicillin-streptomycin (200 IU/L) (Corning) at 37°C, 5% CO_2_, and 100% humidity. Charcoal-stripped fetal bovine serum (Sigma) was used for hormone induced gene expression analysis. SW1736-EV and SW-TRβ cells were modified by lentiviral transduction as recently described^2^. SW1736 and KTC-2 were authenticated by the Vermont Integrative Genomics Resource at UVM using short tandem repeat profiles (SW1736, May 2019; KTC-2, October 2019). 8505C and OCUT2 were authenticated by the U. Colorado using short tandem repeat profiles (8505C, June 2013; OCUT-2, June 2018).

### Pharmacological agents

T_3_ (Sigma-Aldrich) was suspended in 1 M NaOH and GC-1 100% ethanol for stock concentrations of 1 mM before further dilution in culture medium. Buparlisib, Sorafenib, Palbociclib, and Alpelisib (MedChemExpress) were suspended in 100% DMSO for stock concentrations of 50-150 mM before further dilution in culture medium.

### Cell Growth and Viability Assays

Cell growth was measured by cell counting at discrete time points. Cells were seeded in 12- well plates, then treated with GC-1, Buparlisib, Alpelisib, Palbociclib, or Sorafenib to establish time and concentration effects of therapeutics on cell growth. Cell viability was determined by a Sulforhodamine B assay (Abcam) following the manufacturer’s protocol. In brief, ATC cells were plated in 96-well clear flat-bottom plates at a density of 5,000 cells per well. Cells were fixed, stained, and imaged using a plate reader according to manufacturer’s instructions.

### Migration Assay

Cell migration was determined by wound healing assay as previously described^2,23^. Media was supplemented with GC-1 (10 nM) with and without 0.5 μM Buparlisib, 0.5 μM Alpelisib, 1 nM Palbociclib, or 5 μM Sorafenib. Images were obtained at 0, 16, 24, 48, and 72 hours or until 100% wound closure, depending on the cell line. Percent closure was calculated relative to the area of the initial scratch.

### Tumorsphere Assay

Tumorspheres formed from ATC cells were used to assess self-renewal and sphere-forming efficiency as previously described^2^. Where indicated, adherent cells were treated with 0.5 μM Buparlisib, 0.5 μM Alpelisib, 1 nM Palbociclib, 5 μM Sorafenib, or vehicle for 72 hours prior to tumorsphere-forming assay. Tumorspheres were then cultured with or without GC-1 (10 nM) to evaluate the effects of liganded TRβ on thyrosphere growth alone or after treatment with a therapeutic agent.

### Western Blot Analysis

Proteins were isolated from whole cells in lysis buffer and visualized by Western blot as previously described^2,23,25^. Specific proteins were detected with the indicated antibodies (Table 1); immunoreactive proteins were detected by enhanced chemiluminescence (Thermo Fisher Scientific) on a ChemiDoc XRS+ (Bio-Rad Laboratories).

### RNA Extraction and Quantitative Real-Time PCR (qRT-PCR)

Total RNA was extracted using RNeasy Plus Kit (Qiagen) according to manufacturer’s protocol. cDNA was generated using 5X LunaScript RT SuperMix, and mRNA expression was quantified by qRT-PCR using 2X Luna Universal qPCR Master Mix (New England Biolabs) on a QuantStudio 3 real-time PCR system (Applied Biosystems). Fold change in gene expression compared to housekeeping controls was calculated using the ddCT method. Primer sequences are indicated in Table 2.

### Iodide Uptake Assay

Cells were seeded at a density of 40,000 cells per well in a 96 well plate with or without 10 nM GC-1 for 48 hours. Media was discarded, and cells were washed four times with warm PBS before incubation with 1 μM potassium iodide (KI, Sigma-Aldrich) in PBS for 1 hour at 37°C. Wells were washed an additional four times with warm PBS and 100 μL of ddH_2_O were added to each well. 100 μl of 24 mM ammonium cerium (IV) sulfate hydrate with 5.7% concentrated H_2_SO_4_ (Sigma-Aldrich) and 100 μl of 24 mM sodium arsenite (III) with 205 mM sodium chloride (NaCl) and 50 mM sodium hydroxide (NaOH, Sigma-Aldrich) were added to each well. Plates were incubated in the dark at room temperature for 30 minutes, and absorbance at 415 nm was measured with a Synergy 2 Multi-Detection Microplate Reader (Agilent Technologies).

### *In Vivo* Evaluation of Sobetirome (GC-1) and Sorafenib

The xenograft experiment was approved by the Animal Care and Use Committee of the University of Vermont (Protocol X0-018). Four-week-old athymic female nude mice (outbred homozygous nude *Foxn1^nu^/Foxn1^nu^*) were purchased from Jackson Laboratory (Bar Harbor, ME, USA) and were allowed to acclimatize for one week. Mice were given food and water *ad libitum*. Mice were anesthetized using 3% isoflurane delivered at 1.5 L/min, and tumors were established by subcutaneously injecting 1 x 10 8505C cells with a 26G needle in 100 μL of high concentration Matrigel (Corning Inc.) diluted 1:2 with base RPMI-1640 into each flank of 24 mice. We chose 8505C cells based on the rapid growth characteristics *in vitro* observed in our own studies^26^ and based on their ability to form large tumors in *in vivo* studies by other groups^27–30^. One week later, mice were sorted into four treatment groups to achieve approximately equal body weights within each group. Mice were then administered 0.3 mg/kg GC-1, 10 mg/kg Sorafenib, both GC-1 and Sorafenib, or vehicle control every day via intraperitoneal injection using 27G needles. Pharmacological agents were dissolved in 100% DMSO and diluted daily in 30% DMSO, 40% PEG300, and 30% PBS and incubated at 55°C for 10 minutes prior to vortexing to encourage complete solubilization. Tumor dimensions were measured with digital calipers, and the volumes were calculated by the following formula: (**II** x a x b^2^) / 6, where a represents the largest diameter and b is the perpendicular diameter. The body weight of each animal was taken at least twice a week to monitor toxicity. After treatment for 16 days, mice were euthanized with carbon dioxide, and the tumors were harvested, fixed with formalin (Thermo Fisher Scientific), and stored at 4°C prior to slide sectioning and immunohistochemistry.

### Immunofluorescent Analysis of Tumor Tissues

Formalin fixed paraffin embedded human thyroid xenograft tumor blocks were sectioned at 5 µm thickness onto glass microscope slides, dried overnight, and then baked at 60°C for one hour. Sections were deparaffinized in three changes of xylene for 15 minutes each, then dehydrated in a series of graded ethanols (100% x 2, 95% x 2, 70%, 50%, water) for 5 minutes each. Antigen retrieval utilizing 1X DAKO target Retrieval Solution (#S1699, lot# 11270951; Agilent Technologies, Santa Clara, CA) in 50% glycerol was performed in a Decloaking Chamber (SP1 - 100°C for 15 min, SP2 - 90°C for 1 min). Following antigen retrieval, slides were cooled to room temperature in the antigen retrieval solution for 20 minutes. Slides were then rinsed 2x 5 minutes in distilled water followed by 0.2% Tween-20 for an additional 5 minutes. Tissue sections were blocked for non-specific antibody reactivity by incubation with 10% normal goat serum diluted in phosphate buffered saline (PBS)/ 3% bovine serum albumin (BSA)/ 0.3% Triton X-100 for 60 minutes at room temperature. Immediately following blocking, slides were incubated with rabbit polyclonal anti-Ki67 (Abcam, Cambridge, UK; #ab15580) at a concentration 1.4 µg/ml in PBS/3.0% BSA/0.3% Triton X-100 overnight at 4°C. Slides were then rinsed 7x 5 minutes in PBS/3.0% BSA and incubated with goat anti-rabbit Alexa 488 conjugated secondary antibody (ThermoFisher, #A11008) in PBS/3.0% BSA for 60 minutes at room temperature. Staining was followed by 5 x 5 minute rinses with PBS/3.0% BSA. Samples were counterstained with 4’, 6-diamidino-2-phenylindole (DAPI) (ThermoFisher, #D1306) at a concentration of 10 µg/ml in PBS / 3.0%BSA for 15 minutes then rinsed 3x 5 minutes in distilled water. Following rinses slides were mounted with a #1.5 coverslip and DAKO IF mounting media (Agilent; #S3023) for imaging.

### Confocal Imaging of Human Thyroid Xenograft Tissue

Xenograft samples stained with anti-KI-67 and DAPI were imaged on a Nikon A1R HD point scanning confocal microscope with a 20x Plan Apo λ DIC objective lens (NA 0.75, WD 1000 µm) with 488 nm and 405 nm laser excitation respectively. Image acquisition was completed in galvanometic scanning mode (Nikon Instruments Inc., Melville NY). Images of the entire xenograft section were acquired at 12-bit resolution using the tile scan function. Images were saved in ND2 file format in NIS Elements (version 5.2.1, Nikon, Tokyo, Japan).

### Indica Labs HALO Image Analysis

Images were analyzed using HALO image analysis (Indica Labs, Albuquerque NM; version 3.3.45) utilizing the HighPlex FL module (version 4.1.3). Regions of interest containing only tumor tissue were outlined on each tiled image. Nuclear detection was set to the DAPI channel and the nuclear detection settings were optimized to accurately segment nuclei in the images. Membrane and cytoplasmic threshold settings were optimized for Ki-67 staining in the Alexa 488 (green) channel. Cytoplasmic detection settings were set to 4095 (in a 12 bit image) so that no cytoplasmic background staining would be included in the analysis. Nuclear positive signal setting threshold was set per image, to most accurately reflect the antibody staining level observed in the image. Percent Ki-67 positive cells were summarized.

### Ethics Statement and Animal Modeling

All animal procedures were approved by Institutional Animal Care and Use Committee (IACUC) at UVM. All animals were maintained in pathogen-free conditions and cared for in accordance with IACUC policies and certification. All surgeries were performed with isoflurane anesthesia. Temperature-controlled post-surgical monitoring was implemented to minimize suffering. Mice carrying anaplastic thyroid tumors were euthanized at designated time points for tumor collection. We used signs of ulceration or a maximum individual tumor size of 2000 mm^3^ as a protocol-enforced endpoint.

### Statistics

All statistical analyses were performed using GraphPad Prism 9.3.1 and variance assumed similar between experimental groups. Paired comparisons were analyzed by unpaired Student’s t-test assuming two-tailed distribution. Group comparisons were made by one-way ANOVA followed by Sidak’s or Tukey’s multiple comparison test as appropriate. Multigroup analyses was by two-way ANOVA followed by a Tukey’s multiple comparison test. Data are represented as mean ± standard deviation or standard error of the mean where indicated. Differences were considered statistically significant at p□≤□0.05. Area under the curve (AUC) at the 95^th^ confidence interval was used to evaluate statistical differences in growth and migration assays.

## RESULTS

### GC-1 blocks the tumorigenic phenotype of ATC cells transduced with TRβ

We previously demonstrated that T_3_ (10 nM) treatment of transduced SW1736 cells, in which TRβ is re-expressed (SW-TRβ), induced a tumor suppression transcriptomic program and reduced cell growth when compared to cells transduced with empty vector (SW-EV) ^2^. We therefore tested the ability of GC-1 (10 nM) to reduce ATC phenotypes. We found that GC-1 similarly decreased cell growth in four days (**Fig. 1A**). There was a 50% reduction in the overall growth rate of SW-EV cells and 75% reduction in SW-TRβ cells (**Fig. 1B**), indicating potent activation of endogenous TRβ. There was also a modest decrease in the cell migration rate upon treatment with GC-1 (**Fig. 1C**).

**Figure 1.**
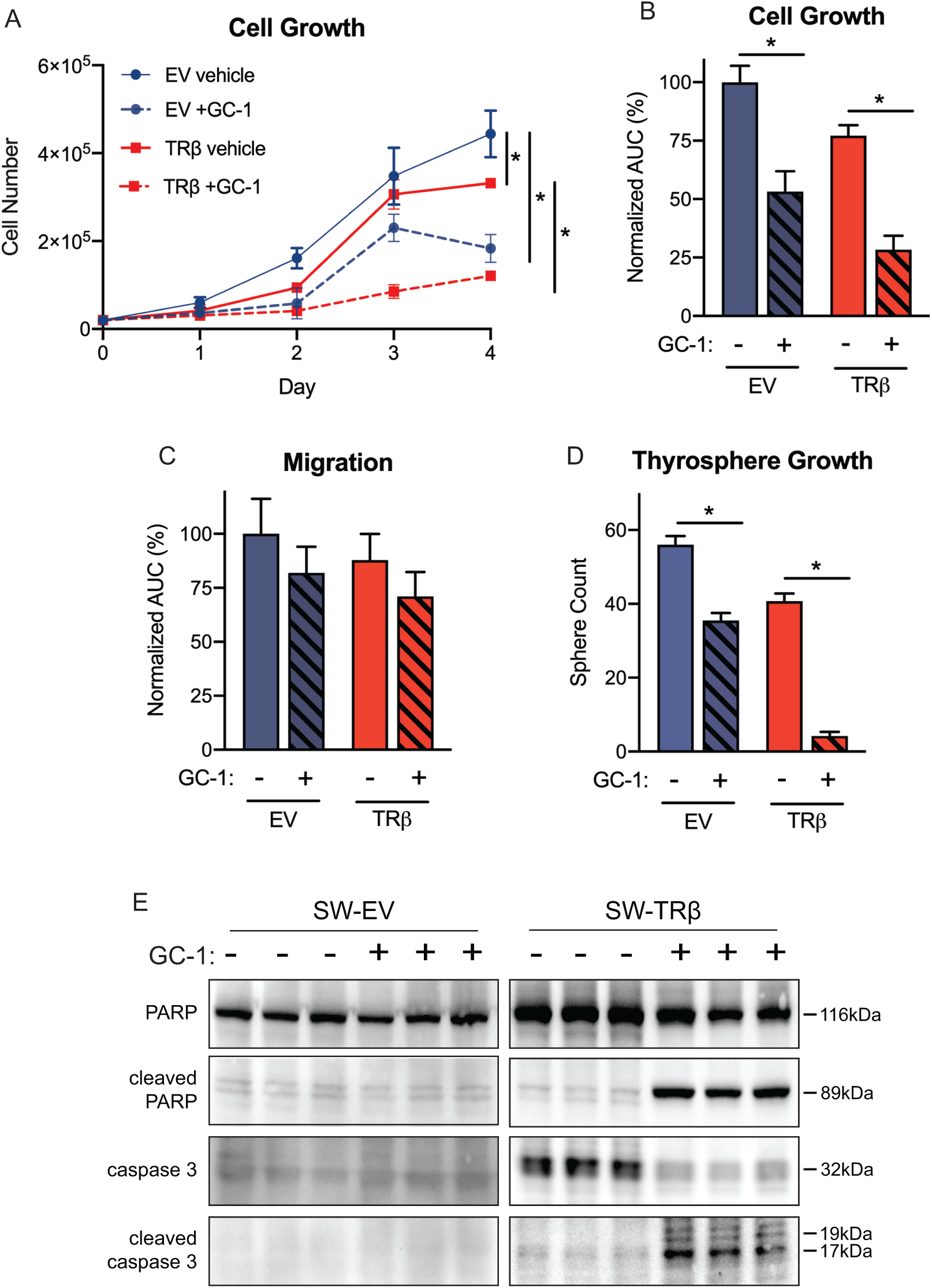
GC-1 blocks tumorigenic phenotypes in ATC cells transduced with TRβ. **A.** Growth curves demonstrate that SW-TRβ growth is significantly decreased after treatment with GC-1 for 4 days (n=3). Significance (* p<0.05) was determined by a two-way ANOVA and Tukey’s multiple comparisons test. **B**. Area under the curve (AUC) analysis of growth curve data. Significance (* p<0.05) was determined by t-test. **C.** Treatment with GC-1 slows migration of SW-TRβ and SW-EV cells measured by scratch assay. Significance (* p<0.05) was determined by t-test. **D.** GC-1 repressed thyrosphere formation in SW-TRβ and SW-EV cells (n=4). Significance (* p<0.05) was determined by t-test. **E.** Immunoblot demonstrates that 5 days of GC-1 treatment induces apoptosis in SW-TRβ cells assessed by cleavage of PARP and caspase 3 (n=3).

As cancer stem cells are thought to be responsible for tumor initiation and recurrence, targeting these cells is key to successful treatment. Therefore, we tested whether thyroid cancer stem cells are responsive to GC-1. Thyrosphere formation is robust in SW-EV cells, and GC-1 induced a significant reduction in growth (**Fig. 1D**). Re-expression of TRβ (SW-TRβ) reduces thyrosphere growth and addition of GC-1 reduces thyrosphere formation by approximately 80% (**Fig. 1D**). These results indicate that GC-1 can activate TRβ to reduce the thyroid cancer stem cell population.

Apoptotic signaling is central to ligand-activated TRβ modulation of the ATC aggressive phenotype ^2^. Thus, we assessed, by immunoblot, the effects of GC-1 on PARP and caspase 3 cleavage as markers of late-stage apoptosis (**Fig. 1E**). GC-1 significantly increased cleavage of PARP and caspase 3 in SW-TRβ consonant with increased apoptotic signaling. Combined, these results indicate that GC-1 can activate TRβ to reduce the aggressive phenotype in transduced ATC cells with TRβ re-expressed and recapitulate effects seen with T_3_ at comparable concentrations.

### GC-1 slows growth of parental ATC cells

Once we confirmed that GC-1 induced the same phenotypic changes as in our transduced ATC cell line model, it was critical to assess whether selective activation of endogenous TRβ could induce similar effects in unmodified ATC cell lines ^24^. SW1736, 8505C, OCUT2, and KTC-2 cells all harbor *BRAF^V600E^* and *TERT* driver mutations, but otherwise have diverse mutational backgrounds (summarized in **Fig. 2A**). Treatment of each cell line with 10 nM GC-1 for four days decreased cell growth whereas treatment with 10 nM T_3_ did not induce a significant change (**Fig. 2B**). Our prior work confirmed that endogenous TRβ expression is low but detectable in the ATC cell lines when compared with normal thyroid cells ^23^. In the present study, GC-1 but not T3 after 24 hr treatment significantly increased TRβ gene expression in SW1736, 8505C, OCUT2 cells (**Fig. 2C)**. Neither GC-1 nor T_3_ treatment of cells significantly altered TRβ protein levels during that time period (**Fig. 2D,E).** Thus, GC-1 but not T_3_ may selectively enhance TRβ gene expression and contribute to stable protein levels in ATC cells.

**Figure 2:**
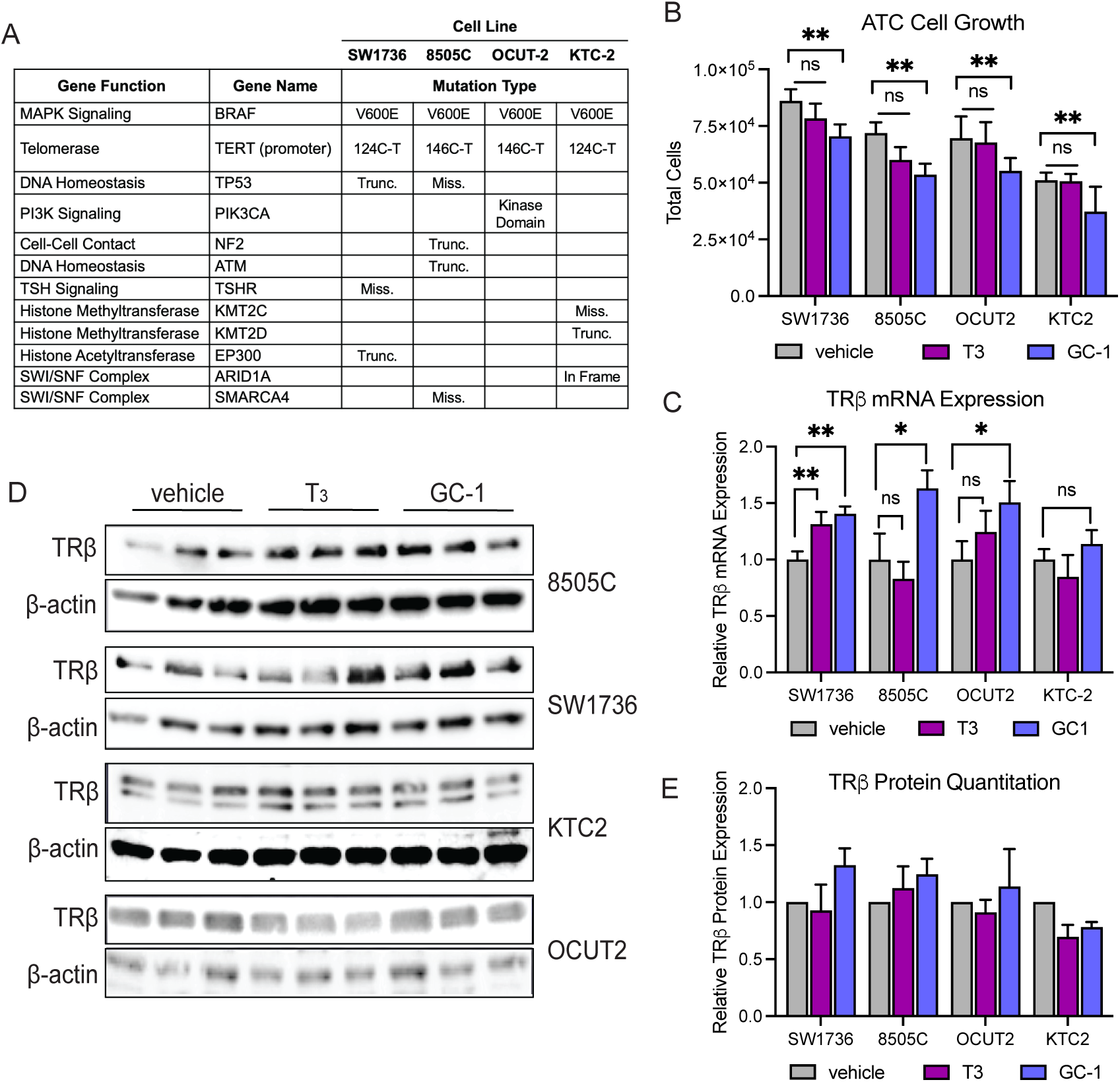
GC-1 slows growth of parental ATC cells. **A**. Summary of mutated genes in four ATC cell lines implemented in this study (17). **B**. Relative cell growth of unmodified ATC cell lines was measured by cell counting assay after 4 days of treatment with 10 nM GC-1 or 10 nM T_3_. ignificance (* p<0.05) was determined by a two-way ANOVA followed by Dunnett’s multiple comparisons test; 3 independent experiments were performed per treatment group. **C.** GC-1 increases TRβ mRNA expression, measured by qPCR, in ATC cells relative to vehicle control at 24 hours. **D**. TRβ protein is detectable by immunoblot in each cell line. **E.** GC-1 treatment induces a modest but not significant increase in TRβ protein levels quantified by densitometry.

### GC-1 increases the efficacy of therapeutic drugs on cell viability

Based on our previous work on activation of TRβ in ATC cells, we hypothesized that GC-1 could enhance the efficacy of therapeutics that target selective pathways. Thus, we chose representative drugs that targeted tumorigenic cell signaling including PI3K, MAPK, and cell-cycle modulated by TRβ. Cell viability was calculated following treatment for three days with increasing concentrations of the PI3K inhibitor Buparlisib (pan-PI3K inhibitor) or Alpelisib (PI3K⍰ mutant inhibitor) for OCUT2 cells which harbor a gain-of-function *PIK3CA* mutation (**Fig. 3A**), the cell cycle inhibitor Palbociclib (**Fig. 3B**), or the multi-tyrosine kinase inhibitor Sorafenib (**Fig. 3C**), with or without 10 nM GC-1. Each therapeutic agent decreased cell viability in three days in a dose-dependent manner. The addition of GC-1 significantly increased the efficacy of these agents at lower concentrations. For example, 1 μM Buparlisib was required to reduce cell viability below 50% in the absence of GC-1, but when combined with 10 nM GC-1 this could be achieved with 0.1-0.5 μM Buparlisib (**Fig. 3A**). Similar trends were observed with all four therapeutics and across the panel of cell lines used.

**Figure 3.**
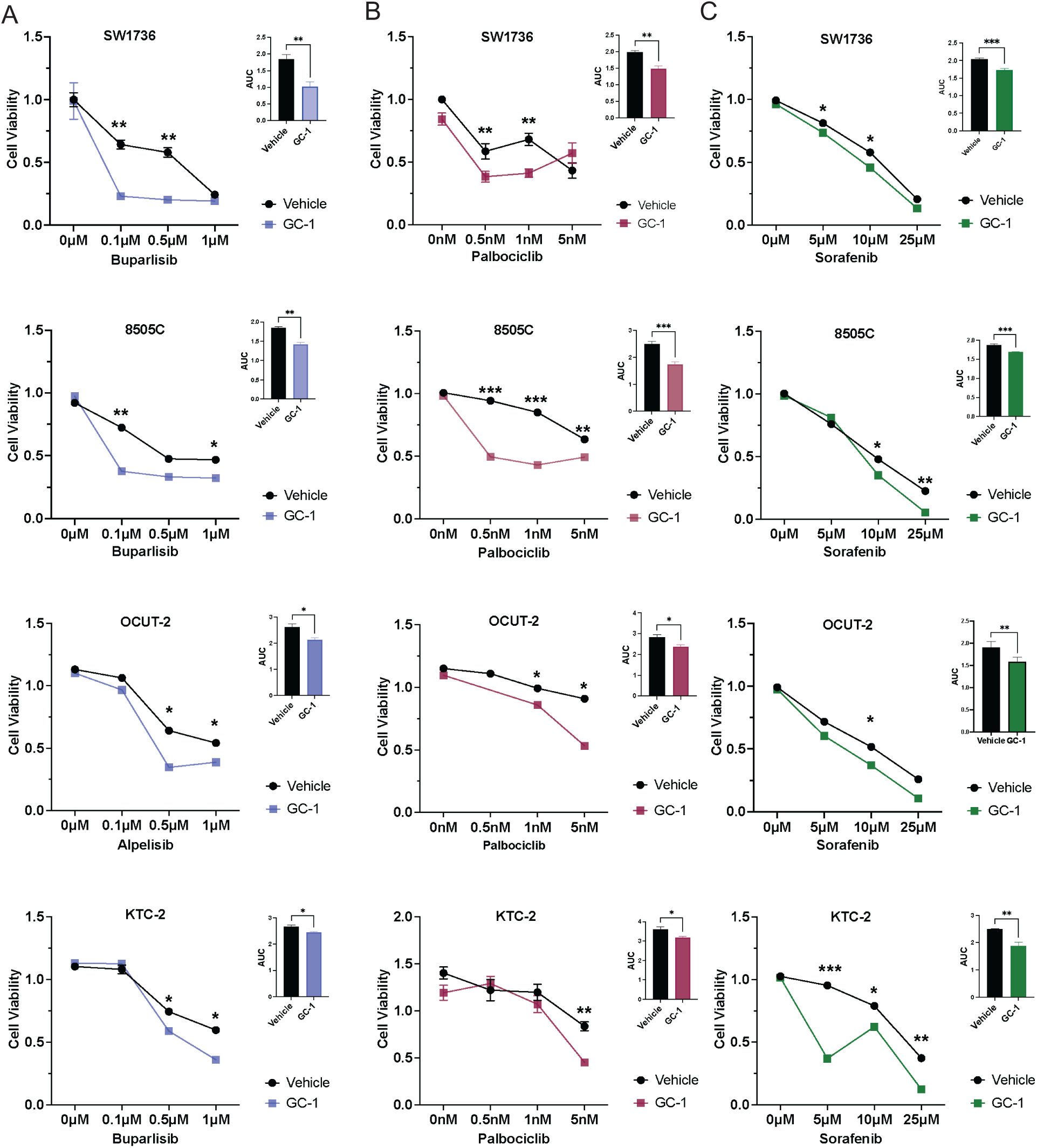
GC-1 increases efficacy of inhibitors. Cell viability of unmodified ATC cell lines was measured by SRB assay after 3 days of treatment with increasing indicated concentrations of buparlisib or alpelisib **A**, sorafenib **B**, or palbociclib **C** simultaneously in combination with 10 nM GC-1. Significance (* p<0.05) was determined by a two-way ANOVA followed by Sidak’s multiple comparisons test; 3 independent experiments were performed per each treatment group. Area under the curve (AUC) analysis of each dose-response experiment is shown in the inset.

### GC-1 enhances the effects of therapeutic agents on cell migration

In addition to reducing cell viability, many therapeutics target pathways that are involved in cell migration. To further evaluate the ability of GC-1 to enhance the efficacy of select therapeutics, cells were treated with the indicated inhibitors at concentrations determined to be effective based on cell viability assays (**Fig. 3**), and migration was observed by assessing wound closure. Each inhibitor was able to successfully slow migration, and this effect was increased when each of the inhibitors were combined with GC-1, except for Buparlisib in KTC-2 cells (**Fig. 4B-E**). Notably, GC-1 alone was able to prevent wound closure of all cell lines with similar success as each inhibitor alone (**Fig. 4A-E**). These findings are similar to what we previously observed in experiments using endogenous thyroid hormone, T_3_, indicating GC-1 is a suitable alternative for TRβ activation.

**Figure 4.**
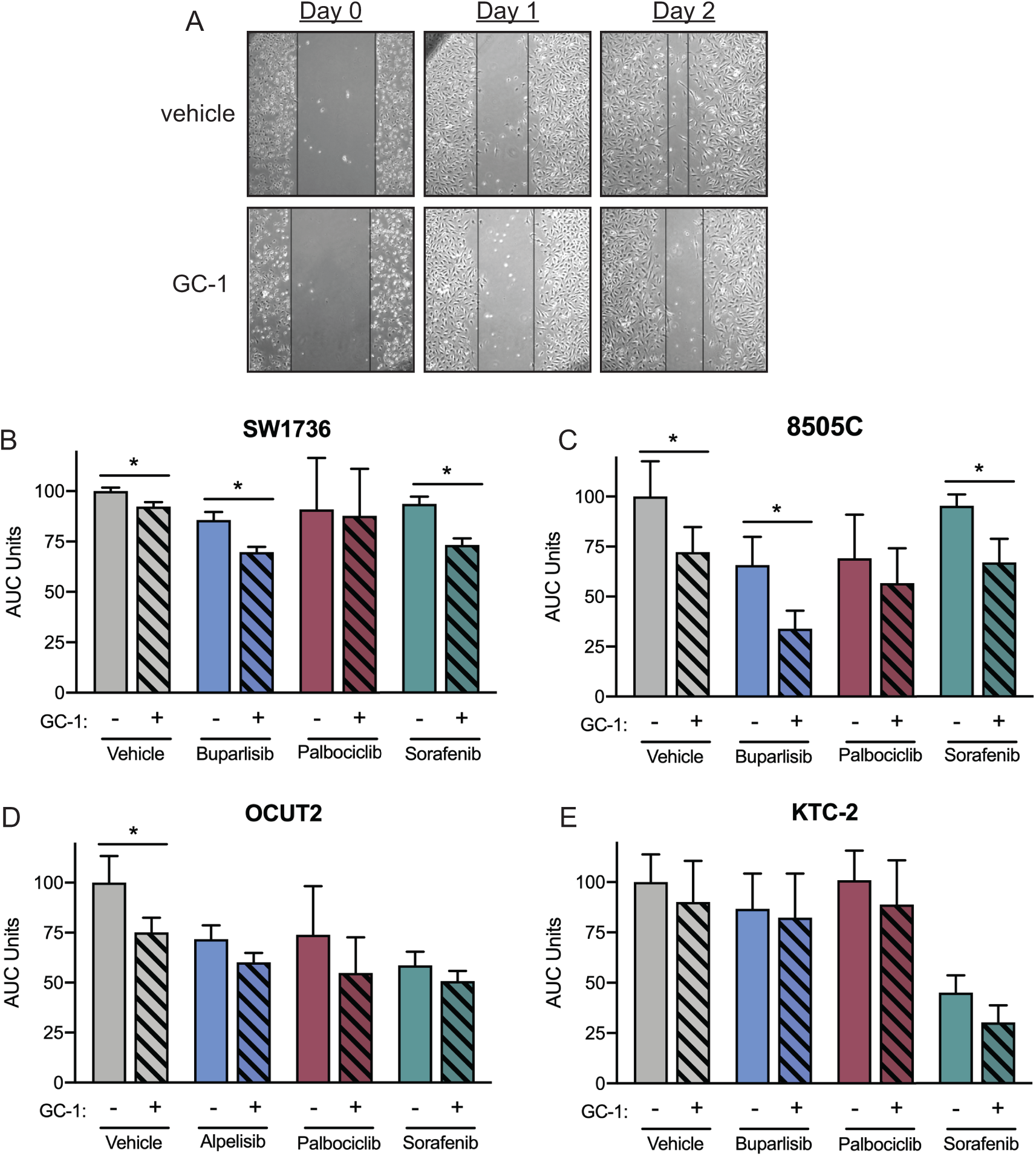
GC-1 enhances the effects of inhibitors on cell migration. Cell migration was measured by scratch assay. Representative images demonstrate reduced scratch closure after two days when ATC cells are treated with GC-1 **A**. Area under the curve (AUC) analysis shows SW1736 **B**, 8505C **C**, OCUT2 **D**, and KTC-2 **E** cells all exhibit lower rates of scratch closer after treatment with GC-1 alone and further reduced scratch closure rates after combined treatment with GC-1 and inhibitors. Significance (* p<0.05) was determined by t-test.

### GC-1 blocks tumorsphere outgrowth and increases efficacy of therapeutic agents

Current treatment options for ATC primarily target hyperproliferative cells, but aggressive cancers like ATC are enriched for long-lived cancer stem cells that are thought to be a major mechanism for therapy resistance and tumor recurrence. We previously demonstrated that ligand-activated TRβ regulates genes related to stemness in ATC cells and that GC-1 could block thyrosphere outgrowth in our modified ATC cells (**Fig. 1D**). We therefore tested the hypothesis that GC-1 could prevent thyrosphere outgrowth after an initial treatment with an inhibitor that targets proliferation.

ATC cells were treated with the indicated inhibitor or vehicle for three days, followed by re-plating in non-adherent conditions. We again observed that GC-1 alone could reduce thyrosphere formation in unmodified ATC cell lines (**Fig. 5A-C**). None of the inhibitors could completely prevent thyrosphere outgrowth when used alone; however, Sorafenib was the most effective overall (**Fig. 5C**). The addition of GC-1 upon removal of each inhibitor blocked nearly all thyrosphere outgrowth. This effect was consistent across all four cell lines and all three inhibitors used. Similar results were obtained with pre-treatment of 8505C cells with GC-1 followed by exposure to therapeutics (**Sup. Fig. 1**). Treatment with GC-1 also decreases mRNA expression of a subset of stemness markers (**Sup. Fig. 2**) This suggests that GC-1 may be particularly effective at blocking cancer stem cell expansion.

**Figure 5.**
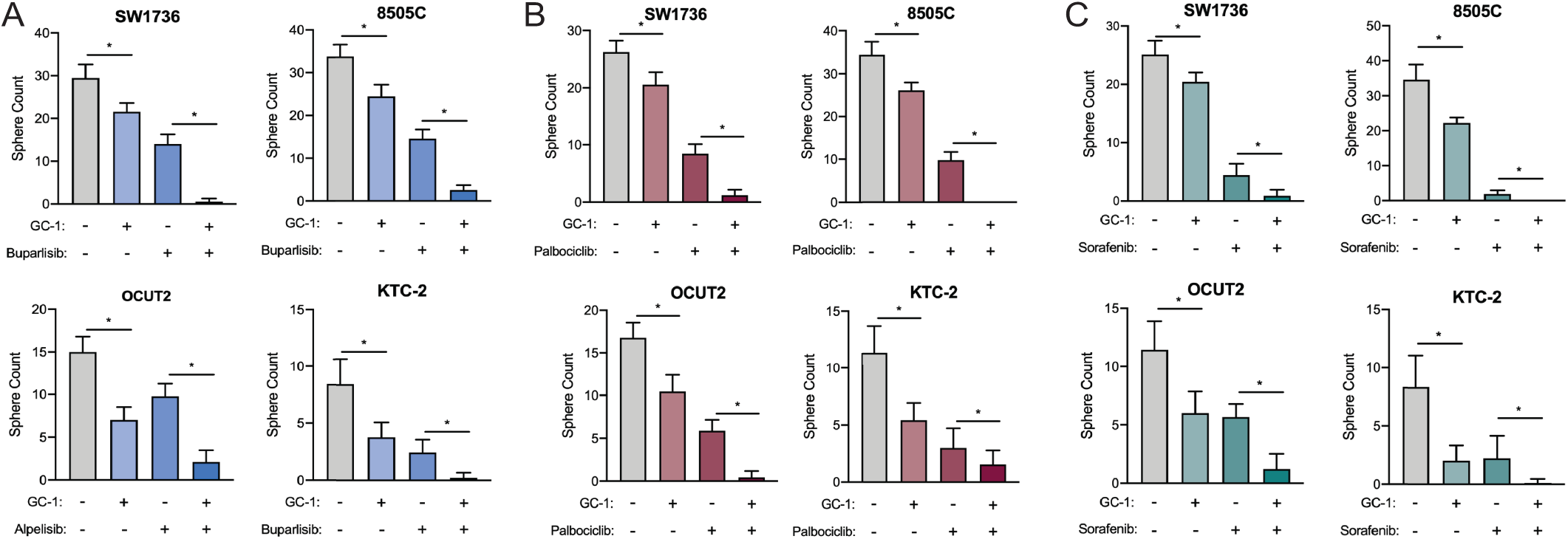
GC-1 blocks thyrosphere outgrowth and increases the efficacy of therapeutic agents. Thyrosphere growth was determined for ATC cells after 3 days of treatment with 0.5 μM buparlisib or alpelisib **A**, 10 μM sorafenib **B**, or 1 nM palbociclib **C** under adherent culture conditions, followed by plating in conditions for spheroid growth in the presence of 10 nM GC-1 for 7 days. GC-1 alone and each therapeutic agent significantly blocked sphere formation in all cell types. GC-1 in addition to each therapeutic agent further inhibited or completely blocked sphere formation. Significance (* p<0.05) was determined by two-way ANOVA followed by Sidak’s multiple comparisons test; 3 independent experiments were performed per each treatment group; n=3 per analyses.

### GC-1 promotes re-differentiation of ATC cell lines

In transduced cells, TRβ activation with T_3_ reduces the aggressive phenotype partially through re-differentiation. Therefore, we investigated the potential for GC-1 to promote re-differentiation in parental SW1736 cells. Gene expression analysis using RT-qPCR revealed that 24 hours of GC-1 exposure increased levels of thyroid-specific gene transcripts (**Fig. 6A**). This transcriptomic program is summarized by the thyroid differentiation score (TDS), which significantly increased in GC-1-treated cells (**Fig. 6B**). Additionally, GC-1 alone decreased the overall number of CD24^-^/CD44^+^ cells present in the monolayer population as evident by RT-qPCR (**Sup. Fig. 3A**). We also observed a decrease in the ATC markers *ALCAM* (CD166) and *ALDH1A1* (**Sup. Fig. 3B**).

**Figure 6.**
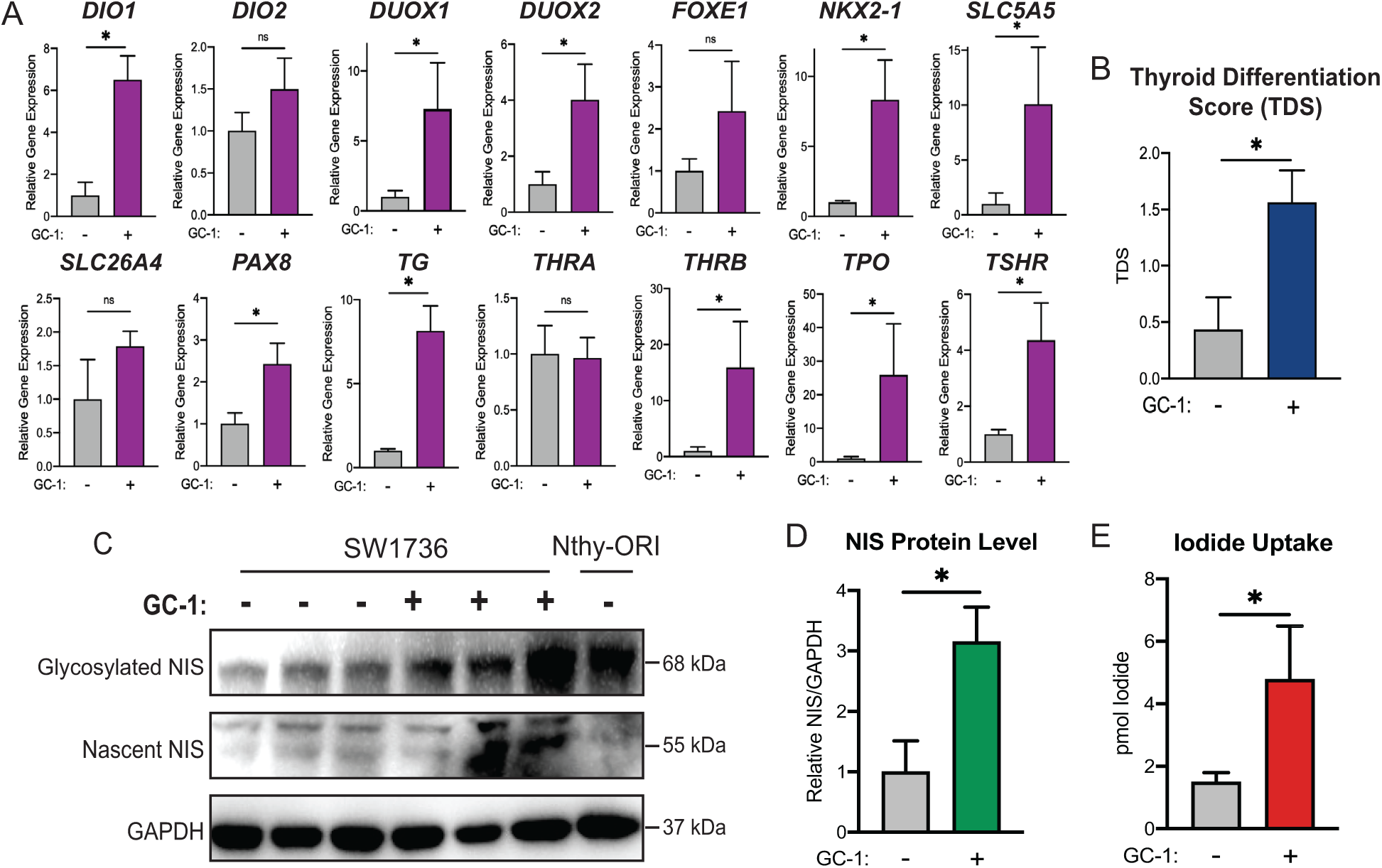
GC-1 induces expression of redifferentiation markers in SW1736 cells. **A.** RT-qPCR for thyroid differentiation markers was performed on SW1736 cells treated with or without 10 nM GC-1 for 24 hours. mRNA levels in cells treated with GC-1 are relative to the untreated cells. Significance (* p<0.05) was determined by Student’s unpaired t test; 3 independent experiments were performed per treatment group. **B.** Thyroid differentiation score (TDS) was calculated using the expression level of the genes in panel A, excluding *THRB*, as previously described (10, 17). Significance (* p<0.0001) was determined by Student’s unpaired t test. **C.** SW1736 cells were treated with or without 10 nM GC-1 in triplicate for 48 hours prior to protein extraction and analysis via immunoblot. Normal-like thyroid Nthy-ORI cells were used as a positive control. **D.** Relative NIS/GAPDH protein levels from C were quantified via ImageJ, and significance (* p<0.01) was determined by Student’s unpaired t test. **E.** Iodide uptake was measured after 48 hours of GC-1 treatment. Significance was determined by Student’s unpaired t test (* p<0.001).

Intriguingly, *SLC5A5* expression was increased with 24 hours of GC-1 (**Fig. 6A**). Therefore, we further investigated NIS protein levels after 48 hours of GC-1 treatment. Both nascent and mature (glycosylated) NIS protein significantly increased following GC-1 treatment (**Fig. 6C-D**). Iodide uptake into cells also increased (**Fig. 6E**), indicating that the increased NIS protein was functional. Similar results were observed in all ATC cell lines including in charcoal stripped serum in 8505C cells to confirm the agonist selective effect (**Sup. Fig. 3**). These findings reveal a novel function of GC-1 to induce re-differentiation in an ATC cell line model and offers a promising target to increase NIS levels and iodide uptake.

### GC-1 blunts tumor growth as effectively as Sorafenib *in vivo*

To validate our cell-based findings *in vivo*, the effect of GC-1 alone or in combination with Sorafenib was tested in a xenograft study. GC-1 blunted tumor growth with no significant change in tumor volume over the treatment period similar to Sorafenib and the combination of both treatments (**Fig. 7A**). At day 16 of treatment, GC- 1 induced a significant 60% reduction in tumor mass as did Sorafenib (**Fig. 7B, C**). The potential for drug synergy was masked due to the equivalent efficacy with Sorafenib at these doses (**Fig. 7A-C**). All mouse groups exhibited similar weight gains over the course of treatment (**Fig. 7D**). Immunofluorescence image analysis (HALO) on excised tumors revealed diminished Ki67 in all treatment groups (**Fig. 7E-F**). The combination of GC-1 and Sorafenib resulted in a significant reduction in Ki67 expression (**Fig. 7F**). Region of Interest (ROI) analysis revealed preferential distribution of Ki67 expression in cells along the periphery of the tumor samples.

**Figure 7.**
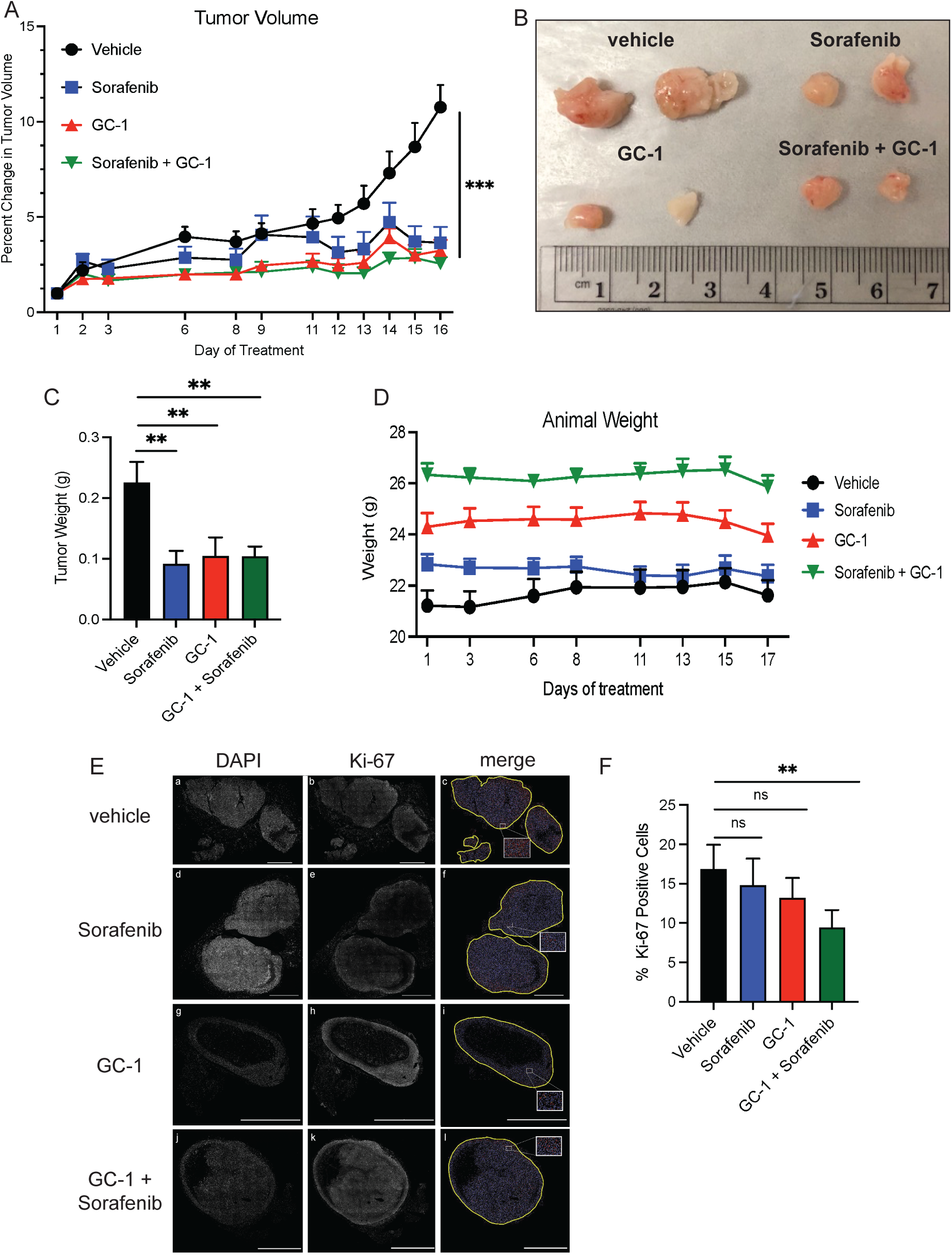
GC-1 inhibits tumor growth as effectively as Sorafenib in an ATC xenograft model. Nude mice were injected subcutaneously with 1 x 10^6^ 8505C cells. Tumors were established after 1 week. Mice were assigned to 4 treatment groups by weight and injected i.p. daily for 16 days with vehicle control, 10 mg/kg sorafenib, 0.3 mg/kg GC-1, or a combination of Sorafenib and GC-1. **A.** Percent change in tumor volumes from day 1 of treatment; *** p<0.001 **B.** Representative excised tumors from each treatment. No evidence of metastases was noted in any group. Reduced vascularization was observed with GC-1 treatment. **C.** Tumors were excised on Day 16 and weighed. Sorafenib (blue), GC-1 (red) induced a 60% reduction in final tumor mass with no difference between the two treatments. The combined treatments did not further reduce the tumor mass over the time period. Significance indicated ** p<0.01. **D.** Mice were weighed three times a week following drug injections to monitor toxicity. No significant change in animal weight was noted for any group. **E.** Representative images of tumors stained for Ki-67 are illustrated. Regions of interest containing only tumor tissue were outlined on each tiled image. Nuclear detection was set to the DAPI channel and the nuclear detection settings were optimized to accurately segment nuclei in the images. **F.** Images were analyzed using HALO image analysis (Indica Labs, Albuquerque NM; version 3.3.45) utilizing the HighPlex FL module (version 4.1.3). Quantification of Ki-67 fluorescent intensity per cell is summarized. Sorafenib and GC-1 decreased Ki-67 but only significantly in combination. Significance (**p<0.01) was determined by one-way ANOVA followed by Sidak’s multiple t-test.

## DISCUSSION

There is compelling evidence that loss of expression of TRβ, a member of the thyroid hormone receptor (TR) family, through genomic modifications and epigenetic silencing is characteristic of thyroid tumors^3,31–33^. Numerous pre-clinical studies demonstrate the potent anti-tumorigenic effect of robust expression of the *THRB* gene. Re-expression of silenced TRβ using demethylating agents delays thyroid tumor progression *in vivo* ^34^, mice expressing a c-terminal frameshift of TRβ (*Thrb^PV^*) spontaneously develop follicular thyroid cancer ^35^, and *Thrb^PV/PV^ Kras^G12D^* mice develop thyroid tumors with features of de-differentiated thyroid cancer ^36^. Our studies in anaplastic thyroid cancer cells revealed that activation of TRβ induces a tumor suppression transcriptomic program, induces re-differentiation, and reduces the aggressive phenotype^2,25^.

Clinical studies of altered thyroidal status have revealed the potential that activation of TR might be of therapeutic value. Patients who were rendered hypothyroxinemic and treated with T_3_ had longer survival with end-stage solid tumors ^37^ and T_3_ treatment enhanced chemosensitivity ^38^. Following tyrosine kinase inhibitor therapy, patients demonstrated improved survival with treatment of hypothyroidism ^39^. Given that TRβ agonists have been developed ^40^, it is remarkable that selective activation of TRβ has not yet been explored in cancer models. Selective agonists have been clinically successful for treatment of metabolic disorders, hyperlipidemia, hypercholesterolemia, and non-alcoholic steatohepatitis without the harmful cardiovascular effects mediated by TRα ^41,42^. In particular Sobetirome (GC-1) and derivatives have been extensively characterized. GC-1 preferentially binds to TRβ and regulates gene expression comparable to T_3_ ^21,43^ and thus a candidate tumor suppressor molecule notably for aggressive cancers such as ATC.

The global incidence of thyroid cancer is increasing faster than any other solid tumor ^44,45^ and outcomes for patients with resistant or recurrent disease and poorly differentiated thyroid cancers (PDTC) are extremely poor ^46^. Patients with advanced or metastatic anaplastic thyroid cancer (ATC) have a higher mortality rate than all other endocrine cancers combined^47^. Thus, novel combinatorial therapies are of critical need.

In ATC tumors, current approaches aim to inhibit or disrupt multiple pathways with combination therapies including blocking cell-cycle, cell survival and proliferation ^48,49^. Our prior work demonstrated that TRβ represses signaling pathways and cell-cycle regulation through genomic mechanisms^2,23^. Therefore, in this study, we tested representative drugs that target the MAPK (Sorafenib), PI3K (Buparlisib, Alpelisib), and cell cycle (Pablociclib) to assess whether GC-1 could increase their effectiveness. GC-1 reduced the effective drug concentrations required to observe a reduction in cell viability. One of our most intriguing observations is that GC-1 alone significantly reduced thyrosphere formation in all ATC cell types. Remarkably, GC-1 in combination with any of the drug treatments nearly eliminated thyrosphere formation.

GC-1 induced re-differentiation in the SW1736 cell line in 24 hours, demonstrating potent transcriptional reprogramming. While we have previously demonstrated the potential for TRβ to promote a more differentiated phenotype in ATC cells, this is the first time we have observed the profound impact of GC-1 in a parental cell line with low TRβ expression. Importantly, we observed a concomitant increase in NIS that led to a higher intake of iodide. Since PDTC and ATC tumors often exhibit a marked decrease in NIS expression and resistance to RAI (radioactive iodine) therapy, the results presented here raise a provocative implication for the use of GC-1 in combination with RAI. Critically, we found that GC-1 could blunt tumor growth as effectively as Sorafenib in a xenograft study confirming the observations in cell-based studies. The combination of GC-1 and Sorafenib did not further reduce the tumor volume or mass over the study period. Future studies establishing the pharmacodynamics of GC-1 in combination with different doses of therapeutic agents will elucidate the most effective combinatorial paradigm.

Our findings demonstrate that selective activation of TRβ with GC-1 can reduce the aggressive tumor phenotype, reduce the cancer stemness in ATC cells, increase the sensitivity of these cells to therapeutic agents, and promote re-differentiation and iodide uptake in transduced cells in which TRβ is re-expressed and in parental cell lines with diverse genetic backgrounds. Moreover, GC-1 treatment is as effective as the therapeutic Sorafenib in inhibiting tumor growth. These studies provide the foundation for further investigation of the broad impact of GC-1 on mitigating tumor progression, pharmaceutical toxicity, and development of resistance to treatment.

## ACKNOWLEDGEMENTS

SW1736 and KTC-2 cell lines were generously provided by Dr. John A. Copland III (Mayo Clinic). 8505C and OCUT-2 cell lines were obtained and authenticated from the U Colorado Cancer Center Tissue Culture Shared Resource supported by National Cancer Institute P30CA046934. Human cell line authentication, NextGen sequencing, automated DNA sequencing was performed in the Vermont Integrative Genomics Resource (RRID# SCR_021775) at UVM. Immunofluorescence staining, imaging and analysis was performed at the Microscopy Imaging Center at UVM (RRID# SCR_018821). Confocal microscopy was performed on a Nikon A1R-HD point scanning confocal microscope (NIH 1S10OD025030-01). We thank the Histology Research Support Facility in the Department of Pathology and Laboratory Medicine at UVMMC. We thank the Office of Animal Care Management for their expertise for *in vivo* experiments.

## Authors’ Contribution Statement

**NEG:** Conceptualization, Formal analysis, Investigation, Writing – Review & editing. **LMC:** Formal analysis, Investigation, Writing. **ERW:** Formal analysis, Investigation. **NMS:** Investigation, Formal analysis. **JAT:** Investigation, Writing – Review & editing. **ELB:** Conceptualization, – Review & editing. **FEC:** Conceptualization, Resources, Writing-Review & editing, Supervision, Project administration, Funding acquisition.

## Funding Statement

The research reported was supported by grants from National Institutes of Health U54 GM115516; National Cancer Institute 1F99CA245796-01; UVM Cancer Center-Lake Champlain Cancer Research Organization (C3) 12577-21; and UVM Larner College of Medicine.

## Authors’ Disclosure Statement

FC is a member of the Endocrinology Editorial Board.

## Data Availability

Original data generated and analyzed during this study are included in this published article or in the data repositories listed in References.

## Notes

### Competing Interest Statement

The authors have declared no competing interest.

### Summary of Updates

This manuscript has been updated with additional data, shown in Figure 2. The text has been revised to reflect the addition of new data.

