## Supplemental Figures for "TRβ Agonism Induces Tumor Suppression and Enhances Drug Efficacy in Anaplastic Thyroid Cancer in Female Mice"

**Supplemental Figure 1**

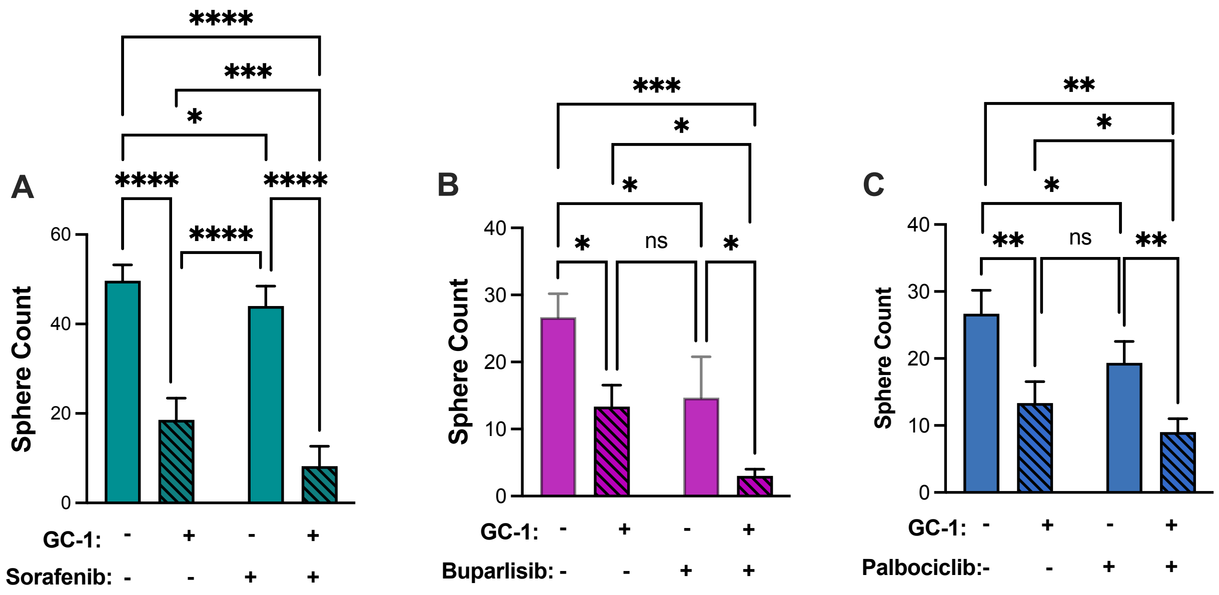


**Supplemental Figure 1.** Pretreatment of 8505c cells with GC-1 blocks thyrosphere outgrowth and increases the efficacy of therapeutic agents. Thyrosphere growth was determined for ATC cells after 3 days of treatment with 10 nM GC-1 under adherent culture conditions followed by plating in conditions for spheroid growth in the presence of 10 μM sorafenib (A), 0.5 μM buparlisib(B), or 1 nM palbociclib (C) for 7 days. GC-1 alone and each therapeutic agent significantly blocked sphere formation in all cell types. GC-1 in addition to each therapeutic agent further inhibited or completely blocked sphere formation. Significance (* p<0.05, ** p<0.01, *** p<0.001) was determined by two-way ANOVA followed by Sidak's multiple comparisons test; 2 independent experiments were performed per each treatment group; n=3 per analyses.

**Supplemental Figure 2**

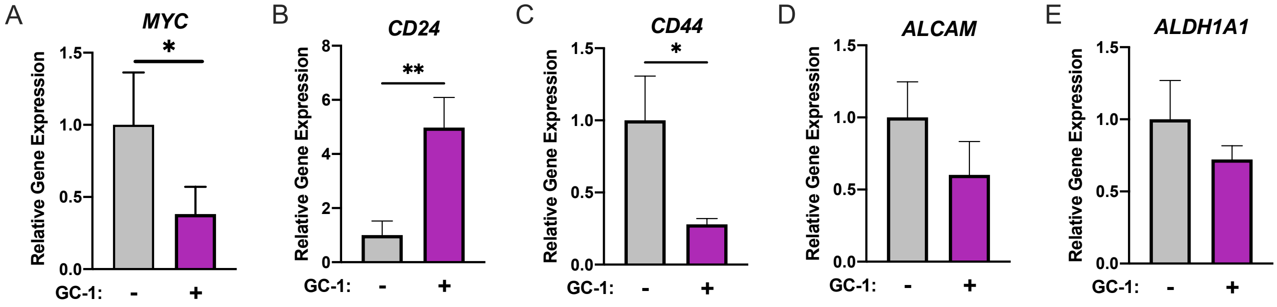


**Supplemental Figure 2.** GC-1 Treatment decreases expression of stemness markers. RT-qPCR for markers of stemness and proliferation was performed on SW1736 cells treated with or without 10 nM GC-1 for 24 hours. mRNA levels in cells treated with GC-1 are relative to the untreated cells. Significance (* $p<0.05$) was determined by Student's unpaired t test; 3 independent experiments were performed per treatment group.

**S. Figure 3**

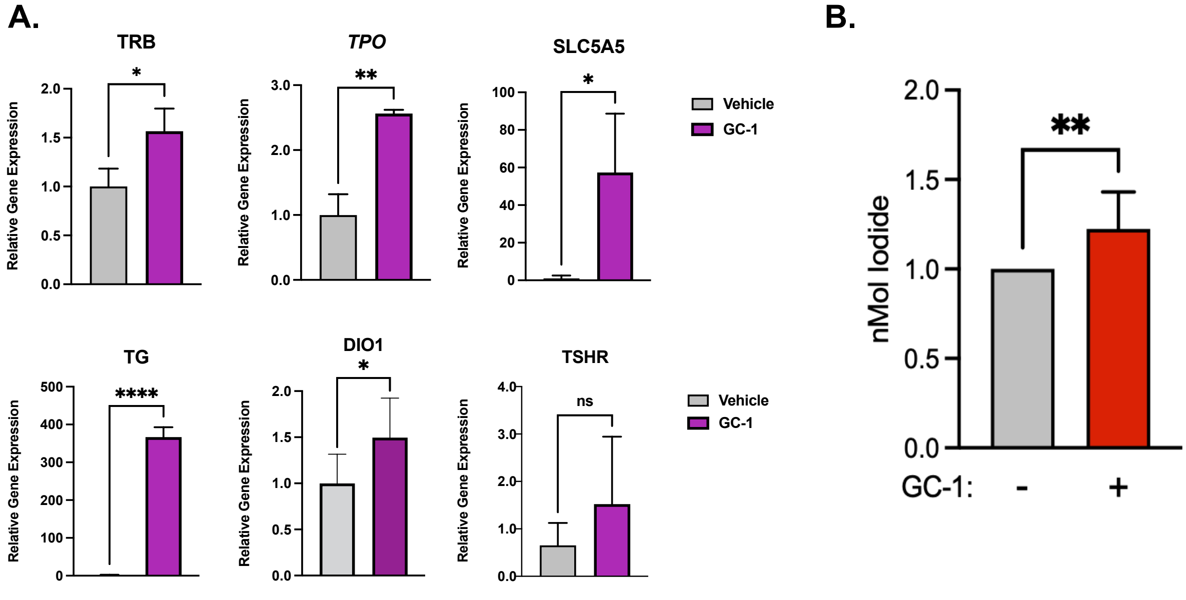


**Supplemental Figure 3. GC-1 induces expression of redifferentiation markers in 8505c cells.** A. RT-qPCR for thyroid differentiation markers was performed on 8505c cells treated with or without 10 nM GC-1 for 24 hours. mRNA levels in cells treated with GC-1 are relative to the untreated cells. Significance (* p<0.05) was determined by Student's unpaired t test, n=3; 2 independent experiments were performed per treatment group. Significance (* p<0.05, ** p<0.01, *** p<0.0001) was determined by Student's unpaired t test. B. GC-1 increases the uptake of iodide in 8505C cells. Iodide uptake was measured after 48 hours of GC-1 treatment. Significance was determined by Student's unpaired t test (* p<0.05), n=32 per treatment group.
