## Supplemental Table 1 for "TRβ Agonism Induces Tumor Suppression and Enhances Drug Efficacy in Anaplastic Thyroid Cancer in Female Mice"

| ***Antibody Table*** | | | | | | | |
| --- | --- | --- | --- | --- | --- | --- | --- |
| **Target** | **Use** | **Antibody** | **Manufacturer** | **Isotype** | **Dilution** | **Antibody Id** | **MW (kDa)** |
| PARP | WB | 9542 | Cell Signaling Technology | rabbit | 1/1,000 | AB_2160739 | 89,116 |
| Caspase-3 | WB | 14220 | Cell Signaling Technology | rabbit IgG | 1/500 | AB_2798429 | 35,19,17 |
| Cleaved Caspase-3 | WB | 9664 | Cell Signaling Technology | rabbit IgG | 1/500 | AB_2070042 | 17,19 |
| NIS | WB | MABC1191 | Sigma Aldrich | mouse | 1/500 | N/A | 75 |
| GAPDH | WB | 600-401-A33 | Rockland Immunochemicals | rabbit | 1/5,000 | AB_2107593 | 37 |
| Goat anti-mouse HRP | WB | 7076 | Cell Signaling Technology | goat | 1/10,000 | AB_330924 |  |
| Mouse anti-rabbit HRP | WB | 211-035-109 | Jackson Immunoresearch Laboratories | mouse | 1/10,000 | AB_2339150 |  |

**Supplemental Table 1**
