## Supplemental Table 2 for "TRβ Agonism Induces Tumor Suppression and Enhances Drug Efficacy in Anaplastic Thyroid Cancer in Female Mice"

| ***Primer Table*** | | | |
| --- | --- | --- | --- |
| **Target** | **Forward (5'-3')** | **Reverse (5'-3')** | **Amplicon Size (bp)** |
| *ALCAM* | TCAAGGTGTTCAAGCAACCA | CTGAAATGCAGTCACCCAAC | 96 |
| *ALDH1A1* | GCACGCCAGACTTACCTGTC | CCTCCTCAGTTGCAGGATTAAAG | 129 |
| *CD24* | TGAAGAACATGTGAGAGGTTTGAC | GAAAACTGAATCTCCATTCCACAA | 208 |
| *CD44* | CCAGAAGGAACAGTGGTTTGGC | ACTGTCCTCTGGGCTTGGTGTT | 151 |
| *DIO1* | CACTGCCTGAGAGGCTCTACATA | TGTAGTTCCAAGGGCCAGAT | 75 |
| *DIO2* | CCTGGTTGCAGCACATTCAC | TTGACTAGCACTGCCTCAGC | 125 |
| *DUOX1* | CCTGGCTCTAGCATGGACAC | TCCCACGAAATGGGGTTCTG | 72 |
| *DUOX2* | GCTGCCTTCCCTTAGTGAGT | TCGCTGGCACTCCATCTTTG | 132 |
| *FOXE1* | CACGGTGGACTTCTACGGG | GGACACGAACCGATCTATCCC | 154 |
| *GAPDH* | ATGTTCGTCATGGGTGTGAA | TGTGGTCATGAGTCCTTCCA | 143 |
| *MYC* | GGCTCCTGGCAAAAGGTCA | CTGCGTAGTTGTGCTGATGT | 119 |
| *NKX2-1* | CTCGCTCATTTGTTGGCGAC | GGAGTCGTGTGCTTTGGACT | 163 |
| *PAX8* | AGTCACCCCAGTCGGATTC | CTGCTCTGTGAGTCAATGCTTA | 139 |
| *SLC5A5* | GCAGTACATTGTAGCCACGAT | TGCAGATAATTCCGGTGGACA | 122 |
| *SLC26A4* | TGAAGGAAATGCCAAAGTTACG | AGTATTCCCGCAGTTTGCTGA | 97 |
| *TG* | AGGGAGAGTTTATGCCTGTCC | CAATACCCAGATACCTCAGGGAA | 148 |
| *THRA* | AGGTCACCAGATGGAAAGCG | AGTGATAACCAGTTGCCTTGTC | 136 |
| *THRB* | CACATCATCATGGTCCAGATGG | GGCGCAGCACGTTGAAAAAT | 92 |
| *TPO* | GCCAACAAGCGGAGTGATTG | GGGCAGCATGTAAGGGAGAC | 175 |
| *TSHR* | TTCCCTGACCTGACCAAAGTT | ACGTCATGTAAGGGTTGTCTGT | 76 |

**Supplemental Table 2**
